# OLIVE Provides Rapid Visualization and Analysis of Chromatin Tracing Experiments

**DOI:** 10.1101/2025.06.18.660347

**Authors:** Yeqiao Zhou, Yimin Sheng, Dongbo Hu, Atishay Jay, Golnaz Vahedi, Robert B. Faryabi

**Author notes:** Equal contributions (Yeqiao Zhou, Yimin Sheng, Dongbo Hu).

## Abstract

Optical chromatin tracing experiments directly capture the three-dimensional folding of thousands of individual alleles, highlighting the need for a tool that enables fast, interactive, and analytical browsing of such data. Here, we introduce Optical Looping Interactive Viewing Engine (OLIVE), a web-based tool designed for high-throughput chromatin tracing data that functions similarly to genome browsers. OLIVE allows users, regardless of computational expertise, to input their own data for automated reconstruction of chromatin fibers at individual alleles or to browse annotated publicly available datasets. Using OLIVE’s functionalities, users can interact with three-dimensional presentation of traced alleles and query them based on spatial features, including pairwise distances and perimeters between their segments. Finally, OLIVE calculates and presents several polymer physics metrics of each allele, providing quantitative summaries for hypothesis-driven studies. OLIVE is an open-source project accessible at https://faryabilab.github.io/chromatin-traces-vis/.

## INTRODUCTION

Chromatin folding has emerged as an important regulator of precise gene expression. Over the past two decades, experimental approaches based on DNA fluorescent in situ hybridization (FISH) imaging have been instrumental in visualizing features of chromatin folding in individual cells ^1^. More recently, Oligopaint DNA FISH probe design and synthesis have greatly increased coverage, resolution, and versatility of DNA FISH-based methods. These advancements, combined with high-throughput imaging, have led to the development of sequential DNA FISH techniques, including Optical Reconstruction of Chromatin Architecture (ORCA)^2^, DNA-MERFISH^3^, DNA seqFISH+^4^, and Hi-M^5^, and Multiplexed Imaging of Nucleome Architectures (MINA)^6^. These methods allow direct tracing of chromatin fiber spatial positioning at thousands of individual alleles at sub-diffraction resolutions, creating a growing need to visualize, assess and interpret results in an intuitive and rapid manner.

To address this unmet need and make chromatin tracing data more accessible to the community, we introduce OLIVE: an easy-to-use, interactive web tool for the reconstruction, visualization and analysis of alleles from chromatin tracing experiments. Using OLIVE’s functionalities, users can automatically render, interact, explore and analyze their own chromatin tracing data. OLIVE also enables users to browse, query and analyze annotated publicly available chromatin tracing datasets.

## RESULTS

Chromatin tracing techniques utilize pools of primary oligos complementary to target genomic sequences, which are linked to barcoded oligos serving as binding sites for fluorescent secondary oligos. Depending on the experimental design, each unique barcode can cover the probes spanning various genomic distances raging from 2 kilobases (Kb) up to 1 megabase (Mb). Chromatin tracing is achieved by sequentially hybridizing, imaging and removing fluorescent secondary probes matching the barcodes one at a time, which are variably referred to as readout, step, or segment.

OLIVE web tool provides a solution for user-friendly and interactive exploration and analysis of thousands of alleles optically mapped by chromatin tracing experiments. Users can upload tabular data containing chromatin segment coordinates based on their experimental configuration or browse through annotated publicly available datasets (Fig. 1A). In both scenarios, users can interact with and query the 3D chromatin fiber reconstruction of individual alleles (Fig. 1B). OLIVE also enables the calculation, visualization and exploration of several distance-based and polymer physics metrics (Fig. 1C).

**Figure 1:**
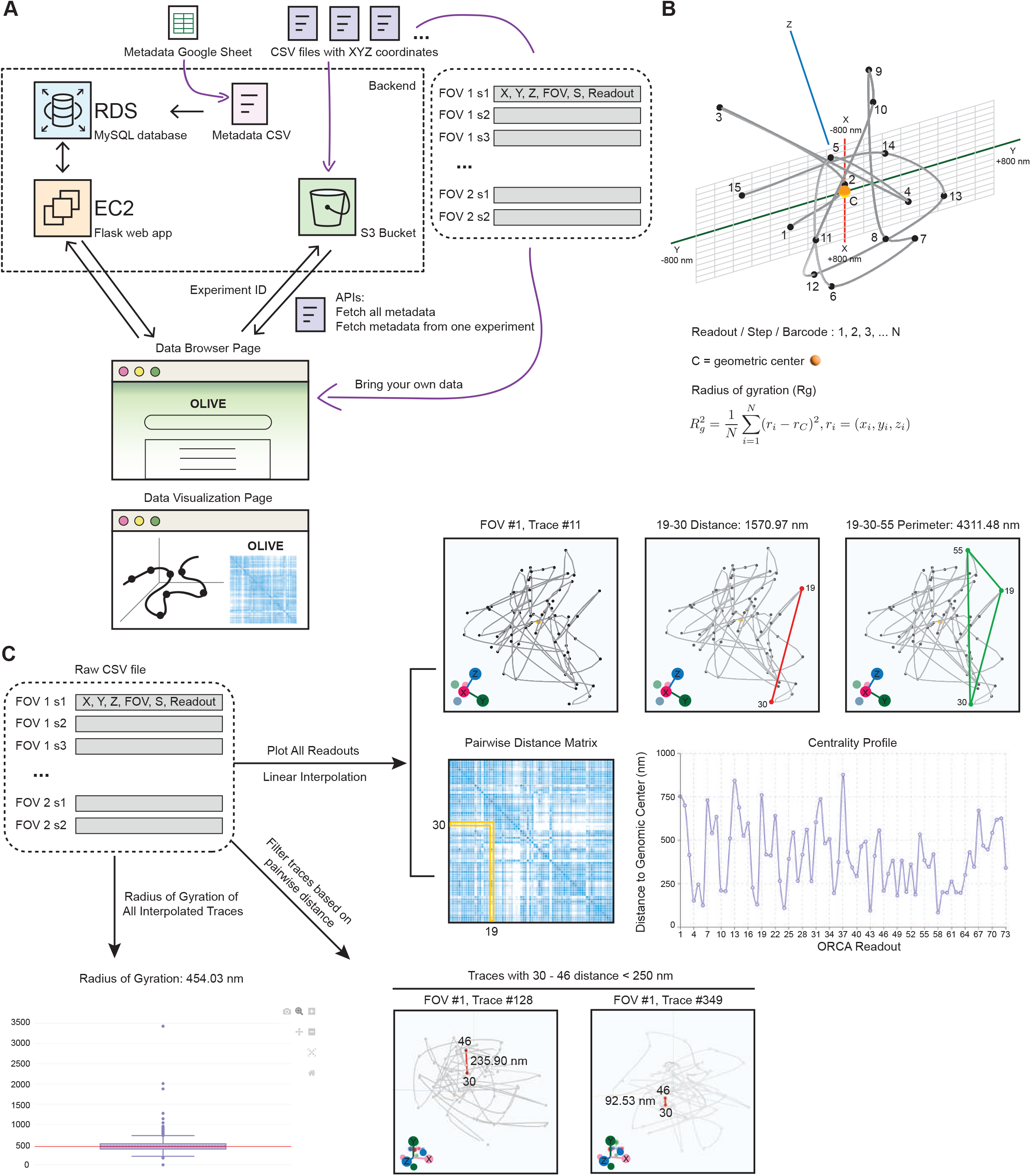
OLIVE enables interactive browsing, query, and analysis of new and publicly available chromatin traces. A: Schematic of OLIVE front end, back end, APIs, web user interface, and data flow. B: Representative view of OLIVE’s chromatin trace visualization (top) and polymer physics metric calculations (bottom). C: Schematic showing possible interactive analysis using OLIVE’s tabular data input. Top-left: required fields in the chromatin trace input file. Top-right: calculation of Euclidian pairwise distances, three-way perimeter, distance matrix, and centrality profile of allele 11 in FOV 1, visualized by OLIVE. Bottom-right: Alleles 128 and 349 were identified using OLIVE’s pairwise distance search, where the distance between readouts 30 and 46 is less than 250 nm. Bottom-left: comparison of the radius of gyration (Rg) of alleles 11 (highlighted in red) with all other alleles in FOV 1 (shown in a box-and-whisker plot).

To support access to published datasets, OLIVE stores data from curated chromatin tracing experiments as an AWS S3 bucket (Fig. 1A). Metadata for each chromatin tracing experiment is maintained in a MySQL database hosted on an AWS RDS instance (Fig. 1A). Communication between OLIVE’s front end and back end database is facilitated through APIs provided by a Flask application running on an AWS EC2 instance. To explore available datasets, users can browse a table, filter and sort columns, and perform global searches to query metadata.

Beyond accessing existing data, OLIVE allows users to upload their own datasets containing coordinates of segments from uniquely identifiable chromatin traces (Fig. 1A). Raw images from different chromatin tracing assays can be processed using tools such as ChrTracer3 and MinaAnalyst, as previously described in detail^7,8^. The output from these programs consists of comprehensive tabular files listing registered alleles along with the X, Y, Z positions of each readout’s optical centroid after corrections for potential aberrations and spot-fitting. OLIVE accepts a simplified version of these outputs, formatted according to 4D Nucleome (4DN) recommendations. For each registered segment of traced chromatins, OLIVE input must include the field of view (FOV), allele number (S), segment (Readout) number, and X, Y, Z coordinates in nanometers, provided in comma-separated format (Fig. 1C). OLIVE provides utility scripts to facilitate these file format conversions.

The folding of individual chromatin traces can be examined in multiple ways. One common approach is to visually inspect reconstructed chromatin by plotting chromatin segment positions as a smoothed 3D curve connecting readouts (Figs. 1B and 1C). Due to experimental challenges – such as target genomic region length, probe specificity, and microscope detection sensitivity – some chromatin traces may have missing readouts. To enable downstream analysis, OLIVE imputes missing readouts using linear interpolation based on flanking readouts. Following imputation, OLIVE reconstructs the 3D structure of each traced chromatin fiber per FOV by first generating a smooth spline tube geometry using the Catmull-Rom algorithm, followed by rendering individual spheres for each coordinate. The geometric center of the trace is then aligned to the origin of the 3D plot. Users can view, zoom, and rotate each trace, which is visualized as a beads-on-a-string 3D object (Fig. 1B).

Using the beads-on-a-string 3D representation of chromatin traces, users can perform a variety of interactive, distance-based analysis in OLIVE (Fig. 1C). Each sphere in the 3D rendering of chromatin trace is a clickable component. When selected, all segments within a specified radius are highlighted. Users can also measure the Euclidean distance between any two selected segments on a chromatin trace. To refine their search for representative alleles, users may filter alleles based on pairwise distances between two selected segments.

Additionally, OLIVE enables users to interactively examine the spatial perimeter formed by any three segments on a chromatin trace – an analysis that informs potential three-way interactions between regulatory elements. OLIVE also automatically computes the Euclidian pairwise distance matrix of each allele and presents the results as an interactive heatmap, linked to its corresponding chromatin trace rendering.

To obtain a more holistic view of chromatin folding, users can compute several polymer physics metrics for each chromatin trace. OLIVE automatically calculates the radius of gyration – a measure of chromatin fiber compaction – for each allele, and compare these values across alleles in the same FOV using a box-and-whisker plot. OLIVE also generates a centrality profile by determining each allele’s geometric center and computing the distance of each segment from that center. Finally, OLIVE offers highly customizable color schemes and output formats, enabling users to download 3D trace rendering and quantitative analysis results as high-resolution, publication-quality images.

## DISCUSSION

Accelerating access to, utilization of, and analysis of chromatin tracing data is critical for advancing single-allele resolution studies of genome folding. To facilitate the exploration and analysis of both new and existing chromatin tracing data, we have developed OLIVE – a unified, interactive web tool that integrated rapid modeling, querying, and quantification of chromatin folding at the allele level. As more datasets are added to OLIVE, the platform will enable the generation of new biological hypotheses and discoveries through rapid and systematic analysis of chromatin tracing data. As of this writing, OLIVE hosts more than 40 annotated chromatin tracing datasets. Leveraging AWS cloud infrastructure, OLIVE is equipped to receive, process, and publish many more. Researchers can submit their chromatin tracing data for inclusion in OLIVE using a streamlined metadata specification form available on the portal. Once published, datasets can be browsed, queried, and analyzed using OLIVE’s computational resources. OLIVE is open-source, well-documented, and accessible at https://faryabilab.github.io/chromatin-traces-vis/ for fast, interactive, and analytical browsing of chromatin tracing data.

## Authors Contributions

Conceptualization: Y.Z., R.B.F.; Methodology: Y.Z., Y.S., D.H., R.B.F.; Investigation: Y.Z., Y.S., A.J., R.B.F.; Resources and Reagents: Y.Z., G.V., R.B.F.; Writing-Original Draft: Y.Z., R.B.F.; Writing-Review & Editing: Y.Z., Y.S., D.H., G.V., R.B.F.; Funding Acquisition: R.B.F., G.V.; Supervision: R.B.F.

## Acknowledgment

The authors thank Shelley Berger for generous support of Penn Epigenetics Institute. This work was supported by NIH grants U01-DK112217, R01-HL145754, R0-AI168240, U01-DK127768 (to G.V.); and R01-CA-248041, R01-CA-230800, U01-DK-112217, U01-DK-123594 (to R.B.F.).

## REFERENCES

1. Shopland, L.S., Lynch, C.R., Peterson, K.A., Thornton, K., Kepper, N., Hase, J., Stein, S., Vincent, S., Molloy, K.R., Kreth, G., et al. (2006). Folding and organization of a contiguous chromosome region according to the gene distribution pattern in primary genomic sequence. J Cell Biol 174, 27–38. 10.1083/jcb.200603083.

2. Mateo, L.J., Murphy, S.E., Hafner, A., Cinquini, I.S., Walker, C.A., and Boettiger, A.N. (2019). Visualizing DNA folding and RNA in embryos at single-cell resolution. Nature 568, 49–54. 10.1038/s41586-019-1035-4.

3. Su, J.H., Zheng, P., Kinrot, S.S., Bintu, B., and Zhuang, X. (2020). Genome-Scale Imaging of the 3D Organization and Transcriptional Activity of Chromatin. Cell 182, 1641–1659 e1626. 10.1016/j.cell.2020.07.032.

4. Takei, Y., Yun, J., Zheng, S., Ollikainen, N., Pierson, N., White, J., Shah, S., Thomassie, J., Suo, S., Eng, C.L., et al. (2021). Integrated spatial genomics reveals global architecture of single nuclei. Nature 590, 344–350. 10.1038/s41586-020-03126-2.

5. Cardozo Gizzi, A.M., Cattoni, D.I., Fiche, J.B., Espinola, S.M., Gurgo, J., Messina, O., Houbron, C., Ogiyama, Y., Papadopoulos, G.L., Cavalli, G., et al. (2019). Microscopy-Based Chromosome Conformation Capture Enables Simultaneous Visualization of Genome Organization and Transcription in Intact Organisms. Mol Cell 74, 212–222 e215. 10.1016/j.molcel.2019.01.011.

6. Liu, M., Lu, Y., Yang, B., Chen, Y., Radda, J.S.D., Hu, M., Katz, S.G., and Wang, S. (2020). Multiplexed imaging of nucleome architectures in single cells of mammalian tissue. Nat Commun 11, 2907. 10.1038/s41467-020-16732-5.

7. Mateo, L.J., Sinnott-Armstrong, N., and Boettiger, A.N. (2021). Tracing DNA paths and RNA profiles in cultured cells and tissues with ORCA. Nat Protoc 16, 1647–1713. 10.1038/s41596-020-00478-x.

8. Liu, M., Yang, B., Hu, M., Radda, J.S.D., Chen, Y., Jin, S., Cheng, Y., and Wang, S. (2021). Chromatin tracing and multiplexed imaging of nucleome architectures (MINA) and RNAs in single mammalian cells and tissue. Nat Protoc 16, 2667–2697. 10.1038/s41596-021-00518-0.

